# Differential Effects of LIS1 On Processive Dynein-Dynactin Motility

**DOI:** 10.1101/124255

**Authors:** Pedro A. Gutierrez, Richard J. McKenney

**Affiliations:** Dept. of Molecular and Cellular Biology, University of California – Davis Davis, CA. USA

## Abstract

Cytoplasmic dynein is the primary minus-end directed microtubule motor protein in cells. LIS1 is a highly conserved dynein regulatory factor that binds directly to the dynein motor domain, uncoupling the enzymatic and mechanical cycles of the motor, and stalling dynein on the microtubule track. Dynactin, another ubiquitous dynein regulatory factor, acts to release dynein from an autoinhibited state, leading to a dramatic increase in fast, processive dynein motility. How these opposing activities are integrated to control dynein motility is unknown. Here we used fluorescence single-molecule microscopy to study the interaction of LIS1 with the processive dynein-dynactin-BicD2N (DDB) complex. Surprisingly, in contrast to the prevailing model for LIS1 function established in the context of dynein alone, we find that binding of LIS1 to DDB does not strongly disrupt processive motility. Motile DDB complexes bind up to two LIS1 dimers, and mutational analysis suggests LIS1 binds directly to the dynein motor domains during DDB movement. Interestingly, LIS1 enhances DDB velocity in a concentration dependent manner, in contrast to observations of LIS1’s effects on the motility of isolated dynein. Thus, LIS1 exerts concentration dependent effects on dynein motility, and can synergize with dynactin to enhance processive movement in the absence of load.

## Introduction

Cytoplasmic dynein (dynein, DYNC1H1) is the only processive minus-end directed MT-based motor protein in the cytoplasm of cells, and as such performs a diverse range of motile activities including transport of small and large vesicular cargos, mRNAs, viruses, and proteins (Allan, 2011; Carter et al., 2016). Dynein is also integral in the construction and positioning of the mitotic spindle (Goshima et al., 2005; Heald et al., 1996; Merdes et al., 1996). In contrast to the single dynein isoform in the cytoplasm, there are ∼15 kinesins that transport various intracellular cargos (Cianfrocco et al., 2015). Specialization of dynein function is thought to arise through distinct dynein subunits, and the interplay between various conserved dynein regulatory factors. Dynein motor activity in cells is predominantly regulated by the LIS1-NudEL complex and the giant ∼1 MDa dynactin complex (Cianfrocco et al., 2015; Kardon and Vale, 2009; Vallee et al., 2012). Mutations in LIS1 and NudEL cause neurodevelopmental diseases while mutations in dynactin cause neurodegenerative diseases, highlighting the diverse roles dynein regulation plays in human neurophysiology (Lipka et al., 2013; Perlson et al., 2010; Vallee and Tsai, 2006).

Haploinsufficency of the *LIS1* gene is the primary cause of the neurodevelopmental disease type 1 lissencephaly (Cardoso et al., 2002; Dobyns et al., 1993). This phenotype is thought to arise from a failure of neuronal precursor cell division and migration, largely due to loss of dynein-driven intracellular nuclear migration (Vallee et al., 2009; Vallee and Tsai, 2006). The lissencephalic phenotype that arises from haploinsufficency of the LIS1 gene suggests that the intracellular concentration of LIS1 is important for proper regulation of dynein motility. Studies of metazoan dynein and LIS1 showed that LIS1 bound directly to the dynein motor domain only when the linker domain was forced into the prepowerstroke conformation (McKenney et al., 2010). LIS1 induced pauses during unloaded dynein motility, and strongly enhanced dynein’s MT affinity when external load was applied via optical trap, leading to greater coordination of force production for teams of dyneins on a single cargo (McKenney et al., 2010). Structural and single molecule studies using yeast proteins confirmed that LIS1 controls dynein’s affinity for the MT track, greatly slowing dynein motility through a direct interaction with the dynein motor domain, near the AAA3/4 interface (Huang et al., 2012). When bound in this position on the motor domain, LIS1 sterically blocked the recovery stroke of dynein’s linker domain, preventing processive movement along the MT (Huang et al., 2012; Toropova et al., 2014). NudEL enhances the interaction of dynein with LIS1 by interacting with the dynein tail domain and recruiting LIS1 to dynein (Huang et al., 2012; McKenney et al., 2010; Zylkiewicz et al., 2011). Thus, the prevailing molecular model from these studies is that LIS1 negatively regulates dynein motility by sterically hindering the remodeling of the linker domain during dynein’s mechanochemical cycle (Cianfrocco et al., 2015; Toropova et al., 2014). This molecular model presents a conundrum for the cellular role of LIS1 function in neurons, where LIS1 is required for dynein-mediated movement of neuronal cell nuclei (Vallee et al., 2009). LIS1’s effect on dynein motility is however beneficial to cargo transport by teams of motors, leading to the idea that LIS1 is crucial for dynein transport of large, high-load intracellular cargos (McKenney et al., 2010; Pandey and Smith, 2011; Yi et al., 2011).

Dynactin was originally isolated as a factor required for dynein transport of vesicles in vitro (Gill et al., 1991; Schroer and Sheetz, 1991), and was reported to modestly enhance dynein’s processivity in vitro (Kardon et al., 2009; King and Schroer, 2000). Dynactin seems to be required for most, if not all, dynein motility in cells and acts as a recruitment, initiation, and processivity factor for dynein motility both in vitro and in vivo (Lloyd et al., 2012; McKenney et al., 2014; McKenney et al., 2016; Moughamian et al., 2013; Nirschl et al., 2016). The CAP-Gly domain of dynactin’s p150*^Glued^* subunit recruits dynactin to dynamic MT plus-ends and is sensitive to the tyrosination state of the MT lattice (Duellberg et al., 2014; McKenney et al., 2016). Point mutations in this domain lead to adult-onset neurodegenerative disease(Levy et al., 2006). Recently, it was discovered that a robust interaction between dynein and dynactin requires the presence of a third, cargo-specific adapter molecule to mediate the interaction between the N-terminus of the dynein heavy chain and the dynactin arp filament (McKenney et al., 2014; Schlager et al., 2014). The formation of this tripartite complex is proposed to force dynein out of an autoinhibited conformation, strongly stimulating processive dynein motility (Urnavicius et al., 2015). Thus, the binding of LIS1 to the motor domain and of dynactin to the tail domain of dynein, have opposing effects on dynein’s mechanochemistry, raising the important question of how the activities of these two regulators are coordinated in the cell for spatiotemporal control of dynein activity.

Here we have examined the interplay between LIS1 and dynactin using purified components in single molecule assays. Surprisingly we readily observe LIS1 binding to motile DDB complexes. Strikingly, LIS1-bound DDB complexes maintained processive movement, in contradiction to predictions made by the current model for LIS1 function (Huang et al., 2012; Toropova et al., 2014). Single LIS1 dimers remain stably bound to DDB over many microns of travel distance, representing hundreds of cycles through the motor’s mechanochemical cycle. Mutations in LIS1 that abolish binding to the dynein motor domain also prevent LIS1 binding to motile DDB. These results suggest that the binding of LIS1 to the motor domain of a dynein molecule that is also activated by dynactin does not inhibit remodeling of the linker domain during dynein’s powerstroke (Toropova et al., 2014). In striking contrast, titration to higher LIS1 concentrations leads to an enhancement of DDB velocity, suggesting that LIS1 may have concentration-dependent regulatory effects on dynein’s mechanochemical cycle. Our results provide new insights into the mechanism of haploinsufficiency found in lissencephalic patients, and suggest that LIS1’s regulatory effects on dynein motility depends on the both the concentration of LIS1 and the presence of other regulatory factors such as dynactin.

## Results

### Single Molecule Observation of LIS1’s Interaction with DDB

To directly observe the effects of LIS1 on DDB motility, we utilized multicolor single molecule TIRF (total internal reflection fluorescence) microscopy. DDB complexes formed with SNAP-TMR-labeled BicD2N (Fig. 1A) moved either processively, or diffused along surface-immobilized taxol MTs with ratios similar to what has been previously described(Hoang et al., 2017; McKenney et al., 2014; Schlager et al., 2014). When we mixed 30 nM of purified, SNAP-647-labeled LIS1 with ∼5 nM DDB, we readily observed LIS1 bound to both processively moving, and diffusing DDB complexes (Fig. 1B, Movie S1). We found that this concentration of LIS1 provided an optimal signal-to-noise ratio in our assays, as higher LIS1 concentrations lead to high background that obscured the LIS1 signal on the MTs, and lower concentrations led to few LIS1 molecules bound to DDB. At this concentration, 18 ± 5 % (n = 612 complexes, two independent trials, SD) of processive DDB complexes contained a bound LIS1 while 34 ± 3 % (n = 320 complexes, two independent trials, SD) of diffusive DDB complexes bound LIS1. It is notable that LIS1 associates with diffusive DDB’s approximately two-fold more frequently than with processively moving complexes under our conditions (P < 0.0001, Fisher’s exact test). We also observed strong accumulation of LIS1 and DDB at the minus-ends of all the MTs (Fig. 1B, Movie S1) (McKenney et al., 2014).

**Figure 1:**
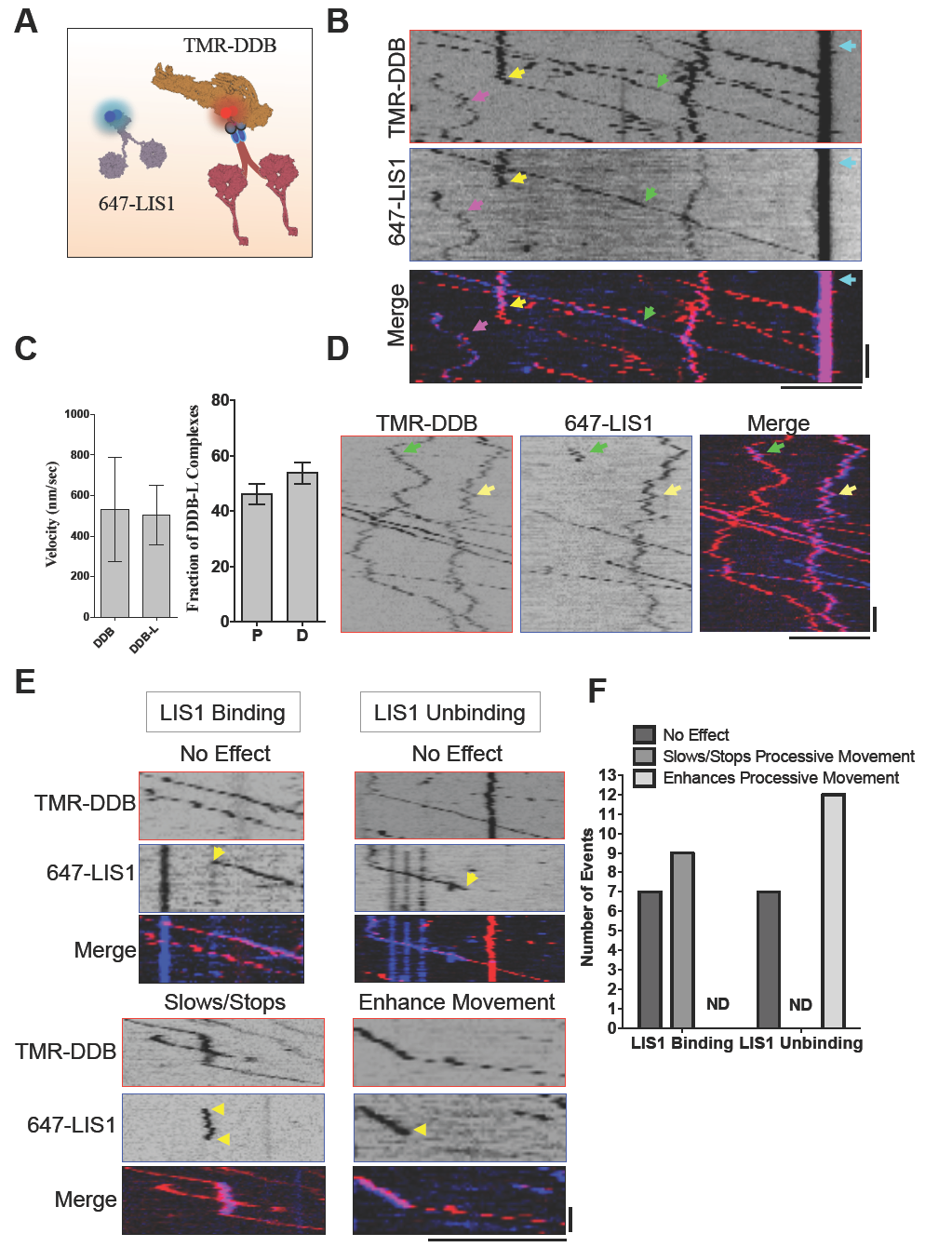
Single Molecule Observation Of LIS1-DDB Interactions. **(A)** Cartoon schematic of the proteins/complexes used in the in vitro reconstitution. LIS1 and DDB are labeled fluorescently via covalent dye conjugation to the SNAP tag on LIS1 and BicD2N respectively. **(B)** Kymographs from individual fluorescent channels showing the movement of SNAP-TMR labeled DDB, and SNAP-647 labeled LIS1. Examples of three types of interactions between LIS1 and DDB are highlighted by arrows: continuous processive motility (green), continuous diffusive motility (pink), and conversion from diffusive to processive motility upon loss of LIS1 signal (yellow). The strong accumulation of DDB and LIS1 signal at MT minus-ends is highlighted as well (teal arrow). Scale bars: 5 μm, 10 sec. **(C)** Quantification of the average velocity of DDB or DDB-L complexes (N = 675 and 435 respectively, 2 independent trials. P = 0.055, unpaired T-test). Error bars S.D. Right, quantification of the fraction of DDB-L complexes that are processive (P) vs. diffusive (D). **(D)** Kymographs from individual fluorescent channels showing examples of transient (green arrow) or stable (yellow arrow) LIS1 binding to diffusing DDBs. Scale bars 5 μm, 10 sec. **(E)** Example kymographs for rare LIS1 binding/unbinding events and their effects on DDB motility. Yellow arrows highlight LIS1 dynamics. Scale bars 5 μm, 5 sec. **(F)** Quantification of observed LIS1 dynamics on DDB motility. Note that LIS1 binding was never observed to enhance velocity, and unbinding was never observed to slow or stop motility (ND, no data).

Surprisingly, the velocity of DDB-LIS1 complexes (DDB-L) was not statistically different from that of DDB complexes lacking a LIS1 signal (Fig. 1C, P = 0.055, unpaired T-test). This is in striking contrast to the effect of LIS1 binding to processively moving yeast dynein, or metazoan dynein lacking dynactin. The ratio of processive to diffusive DDB-L complexes was also similar to that previously observed for DDB complexes in several studies (Fig. 1C) (Hoang et al., 2017; McKenney et al., 2014; Schlager et al., 2014).

The majority of LIS1 remained stably bound to processive DDB during the entire observation period, often for DDB travel distance up to tens of microns (Fig. 1B, Movie S1). We also observed much more rare LIS1 binding or unbinding events (Fig. 1B, D, E, Movie S1). For processively moving DDBs, we classified LIS1 binding/unbinding events as having no effect, enhancing, or retarding processive DDB velocity (Fig. 1D, E). LIS1 binding was approximately as likely to have no discernable effect on DDB motility, as to slow DDB’s velocity (Fig. 1E, F). Dissociation of LIS1 appeared more likely to enhance DDB velocity than to have no observable effect, though the low number of total events precluded statistical analysis (Fig. 1E, F).

We also observed stable LIS1 association with diffusing DDB’s (Fig. 1B, D, Movie S1), but also, albeit more rarely, observed dynamic association/dissociation events (Fig. 1D). We observed that loss of LIS1 signal could correspond to an apparent conversion from diffusive to processive motility (Fig 1B, Movie S1), though this was not always the case (Fig 1D). Binding of LIS1 to a diffusing DDB never caused obvious switching between diffusive and processive motility of DDB, unlike dissociation of LIS1. The activation of motility upon LIS1 dissociation is reminiscent of a LIS1 induced initiation of dynein motility in vivo in *A. nidulans* (Egan et al., 2012). These rare dynamic association/dissociation events provide insight into the multiple effects of LIS1 interactions with the dynein-dynactin complex. We conclude that, when stably bound, LIS1 does not overtly affect DDB motility. However, dynamic association/dissociation of LIS1 can exert control over DDB motility characteristics, suggesting multiple modes of LIS1 regulation of dynein motility.

### LIS1 Binds to the Dynein Motor Domains During Processive DDB Motility

In addition to the previously characterized interaction between LIS1 and the dynein motor domain (Huang et al., 2012; McKenney et al., 2010; Toropova et al., 2014), LIS1 has also been reported to interact with the tail domain of the dynein heavy chain, the intermediate chain of dynein, and the p50 subunit of dynactin (Tai et al., 2002). Because the DDB complex contains all of these potential interaction sites, we therefore set out to map the molecular interaction between DDB and LIS1. LIS1 contains highly conserved residues on one face of its WD40 β-propeller domain (Tarricone et al., 2004) that were previously shown mediate the direct interaction between the *S. cerevisiae* homologues of LIS1 and the dynein motor domain (Toropova et al., 2014). We mutated two of these conserved residues, R316 and W340 in *R. norvegicus* LIS1, to alanine. The purified mutant LIS1 expressed similarly to WT LIS1 and assumed a similar threedimensional shape as WT LIS1 as judged by size-exclusion chromatography (Fig. 2A).

**Figure 2:**
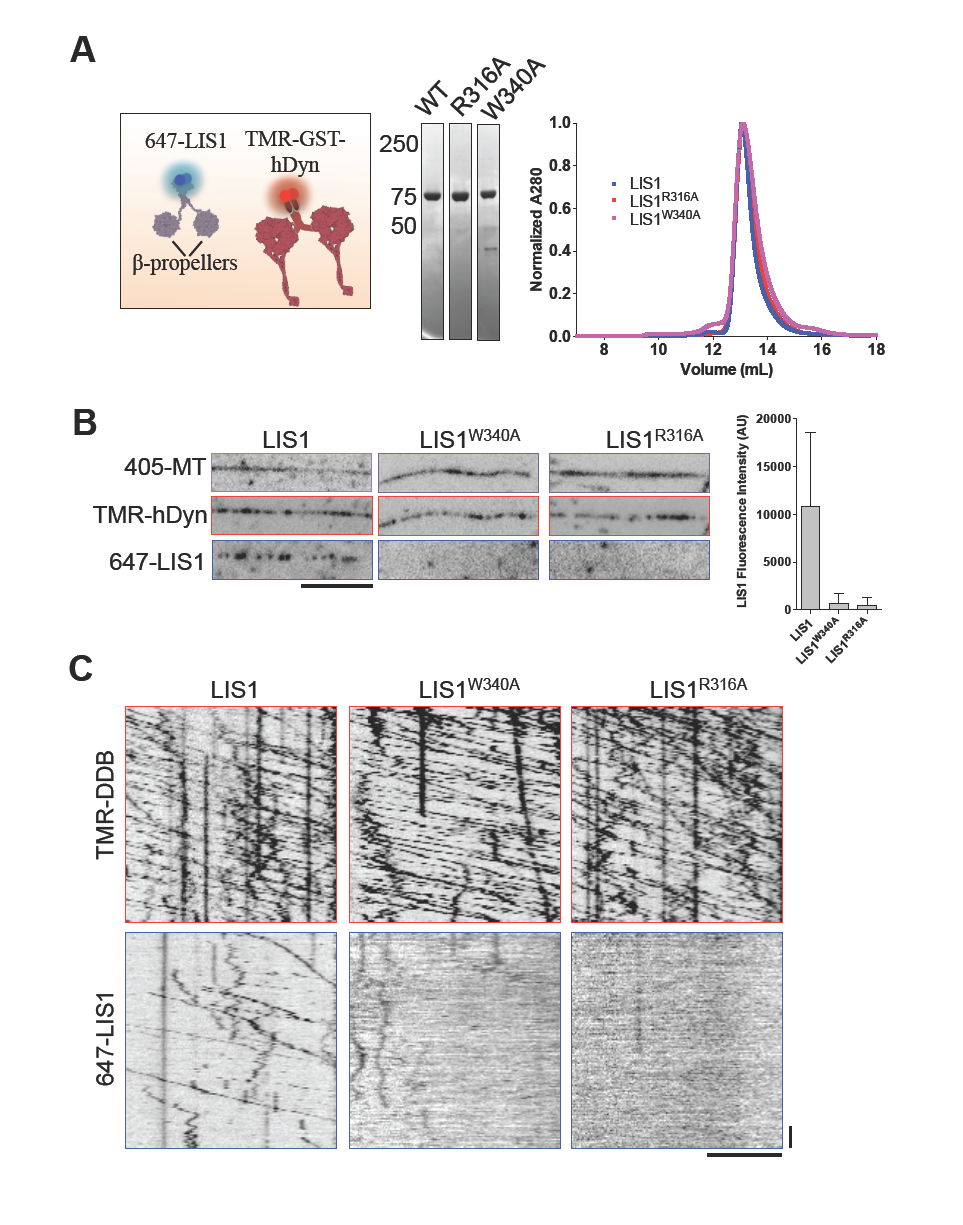
LIS1 Binds Directly To The Dynein Motor Domain During Processive DDB Motility. **(A)** Schematic of the protein constructs used. Note the GST-hDyn contains only the dynein motor domain. Right, Coomassie-stained gel showing recombinant LIS1 constructs used and gel filtration profiles for WT and mutant LIS1 proteins. (**B**) Images from individual fluorescent channels showing Dylight-405 labeled MTs, TMR-SNAP-hDyn, and 647-SNAP-LIS1 signals. Note the mutant LIS1 proteins are not recruited to MTs by SNAP-hDyn.Scale bar 5 μm. Right, quantification of LIS1 fluorescence intensity along MTs for WT and mutant LIS1 molecules (N = 60 MTs quantified for each condition, 2 independent trials). Error bars S.D. (**C**) Example kymographs from each fluorescent channel showing WT 647-LIS1 association with processive TMR-DDB molecules, but no association is observed for either mutant LIS1 constructs, even at high DDB concentrations (∼10 nM DDB, 30 nM labeled LIS1). Scale bars 5 μm, 10 sec.

We next used TIRF microscopy to examine the interaction of LIS1 with recombinant, SNAP-TMR labeled, truncated human dynein (GST-hdyn). In the absence of nucleotide, TMR-GST-hDyn strongly recruited WT LIS1 to surface immobilized MTs (Fig. 2A, B). This is in contrast to the nucleotide-dependent interaction of purified dynein with LIS1 reported previously in solution binding assays(McKenney et al., 2010). This result is particularly notable given that structural data suggest that the linker remains in its unprimed position, sterically incompatible with LIS1 binding (Toropova et al., 2014), when dynein is bound to the MT (Imai et al., 2015). Further experiments to determine the structural conformation of the dynein-LIS1-MT complex will be needed to determine the position of the linker and LIS1 in this co-complex.

We next use this assay to probe the interaction of dynein with mutant LIS1. We observed that neither LIS1^R316A^ nor LIS1^W340A^ were recruited to MTs by GST-hdyn, suggesting both mutants lost the ability to interact with this dynein construct (Fig. 2B). Because GST-hdyn contains only one confirmed LIS1 binding site, we conclude that, similar to the yeast homologues, both LIS1 mutants abolish binding to the dynein motor domain.

We then examined the interaction of the LIS1 mutants with processive DDB complexes. Remarkably, under conditions where we observe robust interaction of WT LIS1 with DDB, we detected no interaction of either mutant with processive DDB complexes, even at high densities of DDB (Fig. 2C). While we cannot rule out that these mutations also affect LIS1’s reported interactions with dynactin and the dynein tail domain, the most parsimonious conclusion from our experiments is that LIS1 binds directly to the dynein motor domain during processive DDB motility.

### Stoichiometry and Stability of LIS1 Binding to DDB

As both LIS1 and dynein are homodimers, it is possible that the stoichiometry of a LIS1-dynein co-complex is 1:1 (dimer:dimer), with each LIS1 β-propeller interacting with a single dynein motor domain. In this configuration, the dynein motor domains would be effectively bridged together through the dimeric LIS1 molecule, possibly restricting the motors conformational space. Alternatively, it is possible that each dynein motor domain binds to one LIS1 homodimer in a 1:2 (dimer:dimer) stoichiometry.

To distinguish these possibilities, we mixed equal molar amounts of two LIS1 populations containing differentially labeled fluorescent SNAP tags with DDB (Fig. 3A). The large majority (91.9 ± 1 %, SD) of processive DDB-L complexes observed only single (either 647-, or 488-SNAP) labeled LIS1 molecules, in approximately equal proportions (647:488 ratio = 1.13 ± 0.27, N = 267 complexes, 2 independent trials SD, Fig. 3B). However, a minority (8 ± 1 %,SD) of DDB-L complexes clearly contained both 647-, and 488-SNAP LIS1 molecules during processive movement (Fig. 3B). In this assay, it is possible that a single DDB complex could be bound by two LIS1 dimers containing the same SNAP label, and thus we conclude that under our conditions, up to 24% of DDB-L complexes contained two LIS1 dimers during processive movement. In addition, we observed rare dynamic events where single DDB-L complexes bound differentially labeled LIS1 molecules at different times during processive, or diffusive, movement (Fig. 3B), further suggesting that most DDB molecules bind single LIS1 dimers in our assay.

**Figure 3:**
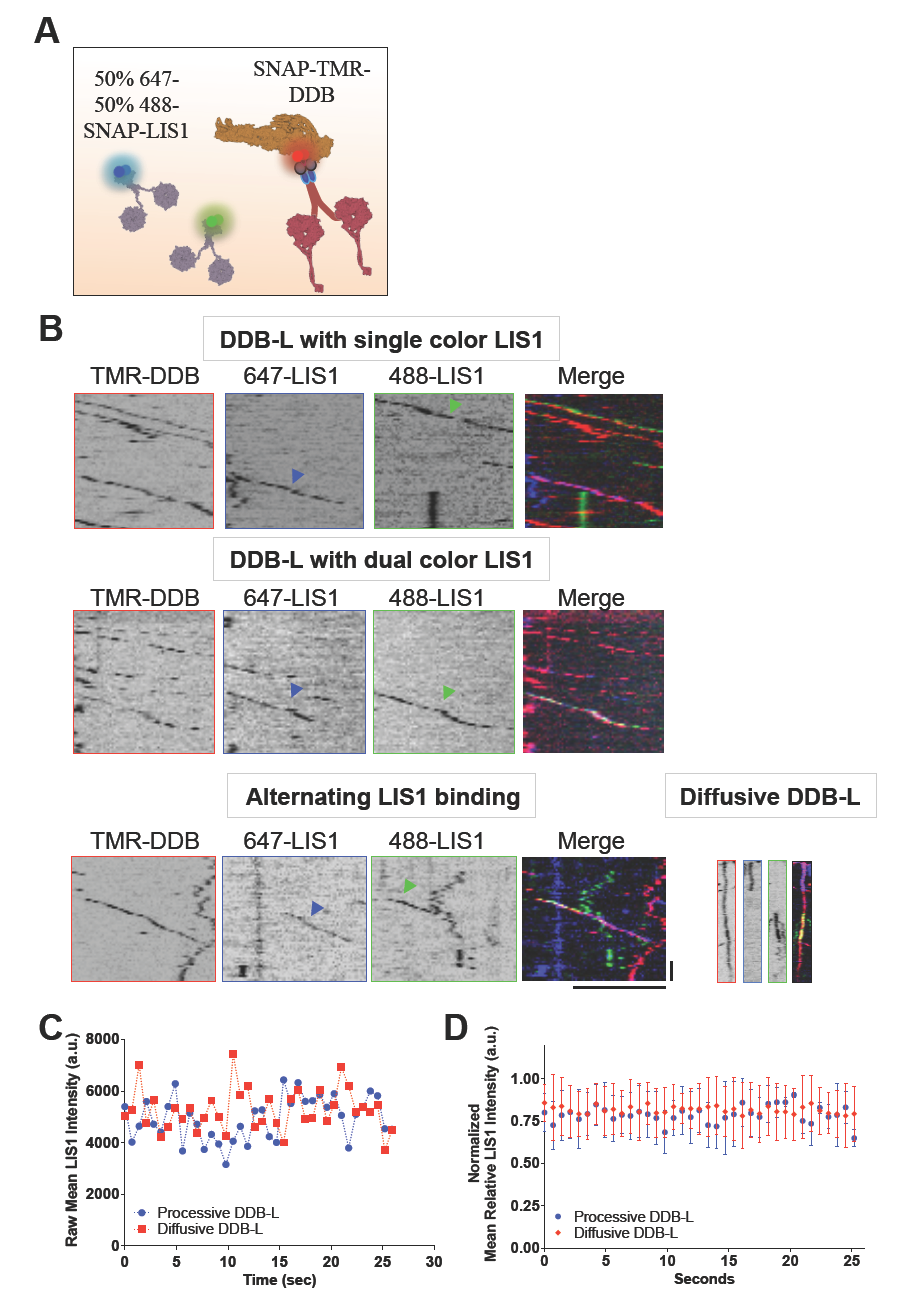
Stoichiometry of Processive DDB-LIS1 Complexes. **(A)** Schematic of the proteins used in the multi-color experiment. **(B)** Kymographs from individual fluorescent channels showing both SNAP-647- and -488 LIS1 molecules (blue and green arrows) and SNAP-TMR-DDB. Examples of individual DDB-L complexes with single color LIS1 (N = 267), or dual-color LIS1 are shown (N = 14, 2 independent trials). Also shown are examples of single processive, or diffusive, DDB complexes alternating between bound differentially labeled LIS1 molecules. **(C)** Example traces from single DDB-L complexes showing the mean LIS1 fluorescent intensity over the entire time-course of the recording. **(D)** Average LIS1 fluorescent signals from processive or diffusive tracked DDB-L complexes were normalized to the highest intensity during the entire trace (N = 10 tracked complexes for each trace, 2 independent trials). Note the fluorescence intensity stays constant during the entire recording suggesting LIS1 is stably bound to DDB.Error bars S.D.

In addition to the experiments described above, we tracked the fluorescence intensity of LIS1 over time for both processive and diffusive DDB-L complexes. Fig. 3C shows typical example intensity traces of individual DDB-L complexes in both categories. The narrow range of LIS1 intensities strongly suggest that the number of fluorescent LIS1 molecules bound to DDB does not change over the course of the track. Normalized averages of many such tracks also show little intensity variance over the observation time (Fig 3D), further suggesting that most LIS1 remains stably bound to DDB. These results suggest that in our experimental conditions, most DDB complexes stably bind a single LIS1 dimer. However, our observations also make it explicitly clear that single DDB complexes are capable of binding two LIS1 dimers simultaneously, and this mode of interaction requires further investigation. This type of interaction may be dependent on LIS1 concentration or other factors that may affect the molecular interaction between LIS1 and dynein, such as post-translational modifications or the presence of other accessory factors such as NudE/L (Kardon and Vale, 2009; Vallee et al., 2012).

### Concentration-Dependent Activation of DDB Velocity by LIS1

The observation of unhindered processive motility even after association of LIS1 with the dynein motor domain challenges the current model for how LIS1 functions to sterically block the dynein powerstroke. At low nanomolar concentrations that facilitate single molecule visualization of this interaction, we do not observe any effect of bound LIS1 on DDB velocity (Fig. 1C).

However, haploinsufficiency of LIS1 causes lissencephaly, strongly suggesting that dynein function may be sensitive to total intracellular LIS1 concentration. To test the effects of LIS1 concentration on DDB motility, we performed single molecule observations of DDB motility in the presence of increasing amounts of LIS1 protein, with LIS1 concentrations much higher than used in the single molecule assays described previously. Strikingly, the addition of up to 1 μM LIS1, approximately a 100-200 fold molar excess over DDB, did not impede processive DDB movement along MTs (Fig. 4A). Addition of 250 nM LIS1 did not significantly affect the overall average velocity of the population, but a four-fold higher amount of LIS1 lead to a statistically significant faster population average (Fig. 4B). We quantified a larger number of processive complexes than in previous reports and noted the resulting DDB velocity distribution histogram was best fit with a sum of two Gaussians (Fig. 4C), showing a distinct shoulder peak at higher velocities. A similar finding was recently reported for the distribution of velocities of dynein-dynactin complexes formed with another adapter protein, Hook3 (Olenick et al., 2016). These observations suggest that activation of dynein by adapter proteins results in a complex distribution of motor velocities, the mechanism of which requires further study.

**Figure 4:**
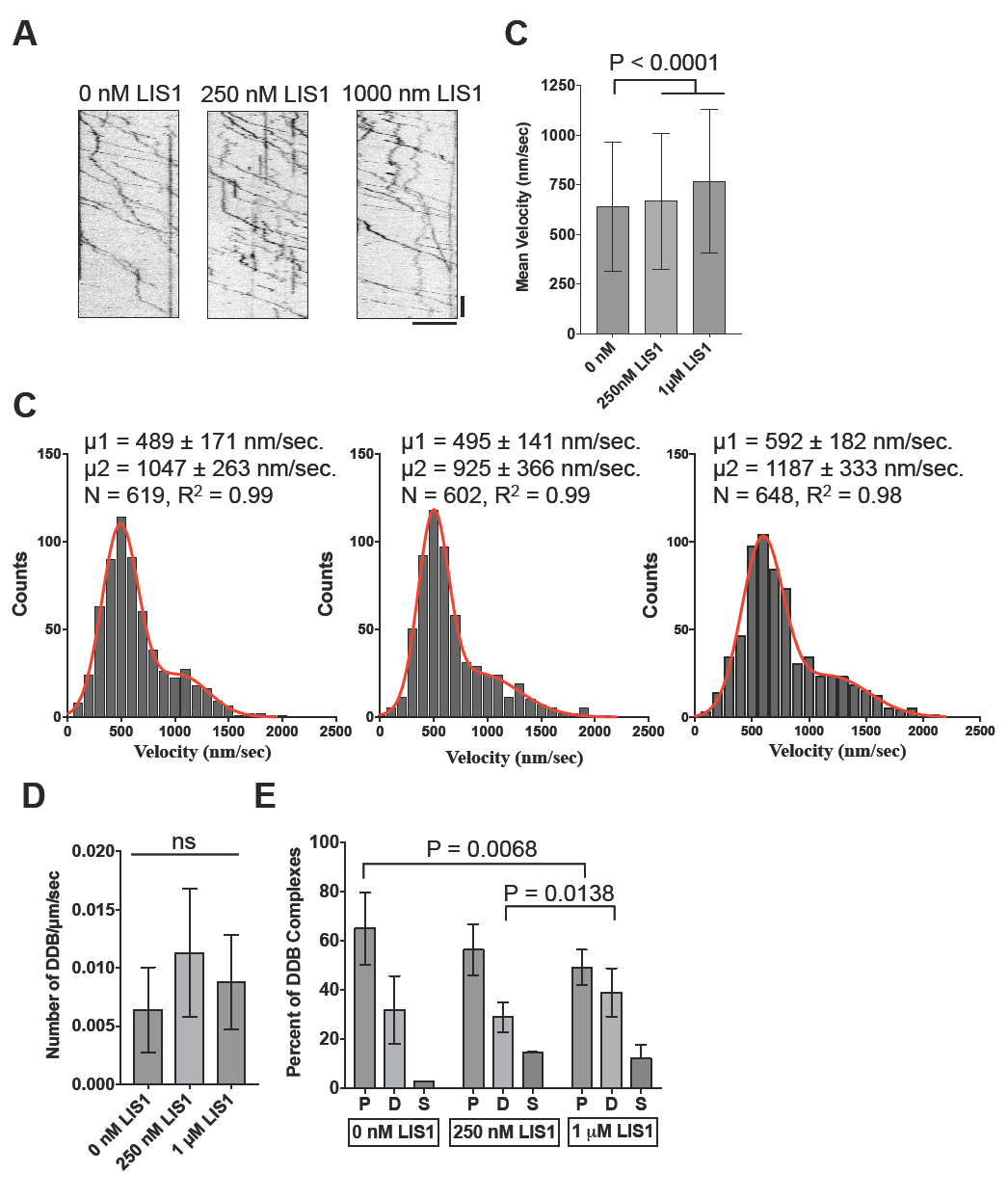
Concentration Dependent Effects of LIS1 On DDB Velocity. **(A)** Example kymographs showing continued processive DDB movement at a range of LIS1 concentrations. Note processive movement continues even at high concentrations of LIS1. Scale bars 5 μm, 10 sec. **(B)** Velocity distribution histograms for each concentration of LIS1 added. Each distribution was best fit by a sum of two Gaussians fit (data are pooled from 3 independent trials). The mean velocities from these fits are shown. **(C)** The population mean for each condition is plotted. Addition 1 μM LIS1 shifts the population mean significantly (P ≤ 0.0001, Kruskal-Wallis test, with post Dunn’s multiple comparison test). **(D)** Plot of the number of DDB complexes per μm of MT per sec. No statistical difference is observed with or without LIS1 (N = 10 MTs and > 200 DDB complexes per condition from at least 2 independent trials). **(E)** Percentage of processive (P), diffusive (D), or static (S) DDB behavior with or without LIS1.

The slower mean velocity within the total population was similar to previously reported values for DDB motility, while the larger mean velocity was nearly two-fold faster (Fig. 4C). The addition of 250 nM LIS1 to DDB did not shift the slower mean velocity significantly (P = 0.5, unpaired T-test), but modestly slowed the faster population (P < 0.0001, unpaired T-test). Surprisingly, the addition of 1 μM LIS1 shifted both populations to higher velocities (P < 0.0001 for both, Fig. 4C, unpaired T-test). These results show that DDB motility is composed of at least two types of velocity parameters, each of which is sensitive to LIS1 concentration.

The amount of LIS1 did not affect the total number of DDB complexes bound to the MT lattice (Fig. 4D). However, LIS1 did shift the percentages of DDB complexes that displayed processive, diffusive, or stationary interactions with the MT lattice (Fig 4E). Increasing LIS1 concentration lead to a decrease of processive DDB complexes by as much as ∼ 24% (P = 0.0068), and increased the number of diffusive and stationary complexes by ∼ 20%, and 75% respectively. Thus high levels of LIS1 facilitate faster DDB velocities at a modest expense of total numbers of processive complexes.

## Discussion

We have used single molecule assays to examine the molecular interactions between two of the most ubiquitous dynein regulatory factors found in cells, dynactin and LIS1. Previous data has suggested that dynactin and LIS1 have opposing effects on dynein motility (Huang et al., 2012; Kardon et al., 2009; King and Schroer, 2000; McKenney et al., 2010; Schlager et al., 2014; Torisawa et al., 2011). Dynactin binding to dynein’s tail domain, mediated through an adapter protein, relieves autoinhibition of the motor and activates fast, extremely processive motility. On the other hand, LIS1 binding directly to the motor domain blocks the structural rearrangement of dynein’s mechanical element, the linker domain, uncoupling ATP hydrolysis from the mechanical cycle, and locking the motor into a high-affinity state on MTs. It is therefore surprising that we observed direct association of single LIS1 molecules with processively moving DDB complexes. A stable association of LIS1 with DDB does not appear to induce dramatic changes to dynein’s mechanochemical cycle, as has been observed in prior studies on dynein in the absence of dynactin (Huang et al., 2012; McKenney et al., 2010; Toropova et al., 2014). However we observed more rare dynamic binding events that do appear to negatively influence DDB velocity, as predicted by the current model for LIS1 function. These results suggest that the effects of LIS1 on dynein motility are multifaceted, and the current understanding of LIS1’s molecular activity (summarized in Cianfrocco et al., 2015) does not encompass the full range of possible LIS1 effects. One possible reason for these observations is that LIS1’s effect on DDB motility may be sensitive to the nucleotide state of the dynein motor domains, including the nucleotide state of AAA3, which gates the MT affinity of dynein and lies near the LIS1 binding site (see below, (Bhabha et al., 2014; DeWitt et al., 2015; Huang et al., 2012; McKenney et al., 2010))

In addition to slowing dynein’s advance along the MT, LIS1 association greatly enhanced dynein’s ability to remain bound to the MT under opposing load in vitro (McKenney et al., 2010), and has been reported to be necessary for the motility of larger, presumably high-load cargos in vivo (Pandey and Smith, 2011; Yi et al., 2011). In addition, isolated lipid droplets were recently shown to retain stably bound dynein, dynactin, and LIS1 proteins. Intriguingly, these droplets dynamically adapt their force output in the face of opposing external load, in a LIS1-depdent manner (Reddy et al., 2016). We cannot determine the effects of opposing load on DDB-L complex motility in our current fluorescence-based assays, but future experiments should focus on exploring the role of load in modulating LIS1’s effects on DDB motility.

How does DDB move processively with LIS1 bound to dynein? Our data with LIS1 mutants suggests that LIS1 is indeed bound directly to the dynein motor domain during DDB motility, likely at the previously mapped interaction site near the AAA3/4 interface (Huang et al., 2012). At this site, LIS1 may play a role in regulating the recently identified AAA3 hydrolysis ‘gate’ that switches dynein between slow and fast motility (DeWitt et al., 2015). The position of LIS1 at this site is sterically incompatible with the swing of dynein’s linker domain during ATP hydrolysis (Toropova et al., 2014).

However, in the DDB complex, it is currently unknown how dynactin binding to the distal tail domain of dynein, which is an extension of the linker, affects the linker position near the dynein motor domain. Conceivably dynactin binding may reposition the linker domain in such a way that allows LIS1 binding near AAA3/4. Intriguingly, tension applied to the linker appears to alter the AAA3 gating mechanism (Nicholas et al., 2015), suggesting a possible mechanism for dynactin-induced changes to linker function. Alternatively, LIS1 has been proposed to enhance the affinity of dynein for dynactin, possibly by relieving an autoinhibited conformation of dynein (Dix et al., 2013; Markus and Lee, 2011). Further work is required to distinguish these possibilities.

Recent cryo-EM data of DDB on MTs (Chowdhury et al., 2015) suggests that the linker domain crosses the dynein motor domain, as previously observed in high-resolution structures of isolated dynein in solution (Carter et al., 2011; Kon et al., 2012; Schmidt et al., 2015). After crossing the motor domain, the linker appears to undergo a pronounced kink, before continuing into the distal tail domain (Chowdhury et al., 2015). We speculate that this kink may provide flexibility to the linker position during DDB motility, and allows LIS1 association. This structure is likely the same as the ‘neck’ region identified in yeast dynein, that when artificially extended, appears to enhance LIS1 association with yeast dynein (Markus and Lee, 2011). Further high-resolution structural work will be required to elucidate the arrangement of the DDB-L complex on MTs.

The concentration dependent effects on DDB motility we observe provide insight into the mechanism of dynein deregulation by loss of a single copy of the LIS1 gene in vivo. At low concentrations in our single molecule assay, stable association of LIS1 does not affect DDB velocity. However, at much higher LIS1 concentrations we observe an activation of DDB velocity. These differential effects could be driven by changes in the stoichiometry of the DDB-L complex. At low concentrations we observe that the majority (75-92 %) of DDB-L complexes bind a single LIS1 dimer, but we also observed binding of two LIS1 dimers to single DDBs, demonstrating that this stoichiometry is possible. We hypothesize that the activation of DDB velocity we observe at high LIS1 concentrations is due to a higher LIS1 occupancy of individual DDBs.

In sum, our data challenge the current molecular model for LIS1 function by demonstrating that LIS1 binding to the dynein motor domain does not sterically impede dynein’s powerstroke when dynein is complexed with an orthogonal regulatory factor, dynactin. Intriguingly, LIS1 forms a tight complex with the dynein regulatory factors, NudE/L (Derewenda et al., 2007; Feng et al., 2000; Niethammer et al., 2000; Sasaki et al., 2000), which have been shown to compete with dynactin for binding to the dynein tail domain (McKenney et al., 2011; Nyarko et al., 2012). NudE/L act to stably recruit LIS1 to dynein (McKenney et al., 2010; Zylkiewicz et al., 2011), and in contrast to LIS1 alone, the NudE/L-LIS1 complex does not induce pausing of processive movement of metazoan dynein along MTs, but retains the ability to induce force-dependent attachment to the MT (McKenney et al., 2010). How the presence of NudE/L affects the ability of LIS1 to interact with DDB will be of particular interest in future experiments.

Thus the effects of allosteric dynein regulatory factors are diverse and dependent on factor concentration and occupancy of individual dynein motors by orthogonal regulators. Our results provide a framework for future experiments to explore how the two most common dynein regulators may simultaneously exert effects on the motors mechanochemistry. How these regulatory factors interact with each other for proper spatiotemporal control of intracellular dynein activity is an outstanding question for future research in vivo, and in vitro reconstitutions such as ours will guide new hypotheses about dynein regulation that can be examined in living cells.

## Movie S1:Example TIRF Movie Showing Complexity of DDB-L Complex Behaviors

Movie shows stably bound, processive DDB-L complex (TMR-DDB in red, 647-LIS1 in blue. Green arrows denote processive complexes) moving tens of microns, a diffusive DDB-L complex (pink arrow), and a complex that switches from diffusive to processive motility upon LIS1 dissociation (yellow arrow, switches to green upon processive movement). Note also strong accumulation of both DDB and LIS1 signals at the ends of MTs. Time is given in seconds and scale bar is 5 μm.

## Acknowledgements

We thank members of the McKenney and Ori-McKenney labs for helpful feedback during this project. We thank Michael Vershinin, Julian Scherer, and Frank McNally for comments on the manuscript.

## Author Contributions

PAG and RJM designed and executed the experiments. PAG and RJM analyzed the data and RJM wrote the manuscript. RJM is funded by an NIH R00 award (4R00NS089428-03).

## Data Availability

The datasets generated during and/or analysed during the current study are available from the corresponding author on reasonable request.

## Materials and Methods

### DNA Constructs

The N-terminal coiled-coil domain of mouse BicD2 (a.a. 25-425), the SNAP-tagged and GST-dimerized human dynein motor domain were previously described (McKenney et al., 2014). *Rattus norvegicus* LIS1 (PAFAH1b1, NP_113951.1) cDNA was obtained from GE Dharmacon and cloned into a modified pFastbacHTA vector containing an N-terminal StrepII-SNAPf tag using Gibson assembly. Mutations in LIS1 were made by site-directed mutagenesis using PCR and Gibson assembly.

### Protein Biochemistry

Porcine brain tubulin was isolated using the high-molarity PIPES procedure as described (Castoldi and Popov, 2003) and then labeled with biotin- or Dylight-405 NHS-ester as described (http://mitchison.hms.harvard.edu/files/mitchisonlab/files/labeling_tubulin_and_quantifying_labeling_stoichiometry.pdf). Microtubules were prepared by incubation of tubulin with 1mM GTP for 10 min. at 37^°^C, followed by dilution into 20 μM taxol for an additional 20 min. Microtubules were pelleted at 80K rpm in a TLA-100 rotor and the pellet was resuspended in 50 μL BRB80 with 10 μM taxol. StrepII-SNAPf-BicD2N was isolated from bacteria as described (McKenney et al., 2014). Purified BicD2N was used to isolate DDB complexes from rat brain cytosol as previously described (McKenney et al., 2014). DDB complexes were labeled with 5 μM SNAP-TMR dye during the isolation procedure, and were frozen in small aliquots and stored at −80^°^C.

Baculoviruses for LIS1 constructs were made according to the Bac-to-Bac protocol using SF9 cells (Invitrogen). SF9 cells were infected at 1-2×10^6^/mL and typically P2 virus was used for protein expression at a 1:100 dilution. Infections were allowed to proceed for ∼60 hrs and cells were harvested and the cell pellet was frozen and stored at −80^°^c. Cells were thawed and resuspended in lysis buffer (50mM Tris, PH 8.0, 150mM K-acetate, 2mM MgCl2, 1mM EGTA, 10% glycerol, 1mM PMSF, 1mM DTT). Cells were lysed by incubation with 1% Triton X-100 for 15 min on ice and 5 passages through a dounce homogenizer. The cell pellet was centrifuged at 16,000 × g, and the supernatant was passed over a column packed with Streptactin Superflow resin (Qiagen). The column was washed with four column volumes of lysis buffer and bound proteins were eluted in lysis buffer containing 3mM desthiobiotin (Sigma). Eluted proteins were concentrated on Amicon filters and passed through an EnRich650 (Bio-Rad), or superpose 6 (GE Healthcare)gel-filtration column, in lysis buffer, using a Bio-Rad NGC system. Peak fractions were collected and again concentrated and frozen in small working aliquots. For labeling with SNAP dyes, proteins were labeled with 2-5X molar excess of SNAP dye (SNAP-Alexa 647 or SNAP-Surface 488) for 2-4 hrs on ice. The unbound dye was removed using Zebaspin columns (Invitrogen). The stoichiometry of labeling was assessed using a Nanodrop One (ThermoFisher) and comparing the absorbance of total protein at 280nm to the absorbance at the SNAP-dye wavelength. For TIRF experiments, we only used preparations where labeling of protein was greater than 60%. Concentrations given are for the total labeled amount of protein in the assay and are 24 calculated for the monomeric forms of the proteins. All buffers and chemicals were from Sigma.

### TIRF Microscopy and Data Analysis

All microscopy was performed on a custom built through the objective TIRF microscope (Technical Instruments, Burlingame CA) based on a Nikon Ti-E stand, motorized ASI stage, quad-band filter cube (Chroma), Andor laser launch (100 mW 405 nm, 150 mW 488 nm, 100 mW 560 nm, 100 mW 642 nm), EMCCD camera (iXon Ultra 897), and high-speed filter wheel (Finger Lakes Instruments). All imaging was done using a 100X 1.45NA objective (Nikon) and the 1.5X tube lens setting on the Ti-E. Experiments were conducted at room temperature. The microscope was controlled with Micro-manager software (Edelstein et al., 2010).

TIRF chambers were assembled from acid washed coverslips (http://labs.bio.unc.edu/Salmon/protocolscoverslippreps.html) and double-sided sticky tape. Taxol-stabilized MTs were assembled with incorporation of ∼ 10% Dylight-405-, and biotin-labeled tubulin. Chambers were first incubated with 0.5 mg/mL PLL-PEG-Biotin (Surface Solutions) for 10 min., followed by 0.5 mg/mL streptavidin for 5 min. Unbound streptavidin was washed away with 40 μL of BC buffer (80mM Pipes pH 6.8, 1mM MgCl_2_, 1mM EGTA, 1 mg/mL BSA, 1mg/mL casein, 10μMtaxol). MTs diluted into BC buffer were then incubated in the chamber and allowed to adhere to the streptavidin-coated surface. Unbound MTs were washed away with TIRF assay buffer (60 mM Hepes pH 7.4, 50 mM K-acetate, 2 mM MgCl_2_, 1 mM EGTA, 10 % glycerol, 0.5 % Pluronic F-127, 0.1 mg/mL Biotin-BSA, 0.2 mg/mL κ-casein, 10μM taxol). Typically ∼ 5 nM DDB and 30 nM labeled LIS1 was then diluted into TIRF assay buffer containing 2 mM Mg-ATP and introduced into the chamber. Images were acquired every ∼ 0.5-0.7 seconds. For binding to GST-hDyn, ∼10 nM purified and TMR-labeled motor was incubated with 0.1 unit/mL apyrase, to induce rigor binding to the MTs, and 30 nM each LIS1 species in the TIRF chamber for 10 minutes before images were acquired.

The resulting data was analyzed manually using kymograph analysis in imageJ (FIJI). For velocity analysis, the velocity of an uninterrupted run segment from a kymograph was used. Approximately 10-20 % of the DDB complexes paused or changed velocity during the run and the velocity of the initial run segment before the change was used. The TrackMate plugin was used to analyze mean fluorescence intensity over time (Fig. 3C-D). For images displayed in figures, background was subtracted in FIJI using the ‘subtract background’ function with a rolling ball radius of 25 and brightness and contrast settings were modified linearly. Graphs were created using Graphpad Prism 7.0a and statistical tests were performed using this program. All variances given represent standard deviations.

## References

Allan, V.J. 2011. Cytoplasmic dynein. Biochem Soc Trans. 39: 1169–1178.

Bhabha, G., H.C. Cheng, N. Zhang, A. Moeller, M. Liao, J.A. Speir, Y. Cheng, and R.D. Vale. 2014. Allosteric communication in the dynein motor domain. Cell. 159: 857–868.

Cardoso, C., R.J. Leventer, J.J. Dowling, H.L. Ward, J. Chung, K.S. Petras, J.A. Roseberry, A.M. Weiss, S. Das, C.L. Martin, D.T. Pilz, W.B. Dobyns, and D.H. Ledbetter. 2002. Clinical and molecular basis of classical lissencephaly: Mutations in the LIS1 gene (PAFAH1B1). Human mutation. 19: 4–15.

Carter, A.P., C. Cho, L. Jin, and R.D. Vale. 2011. Crystal structure of the dynein motor domain. Science. 331: 1159–1165.

Carter, A.P., A.G. Diamant, and L. Urnavicius. 2016. How dynein and dynactin transport cargos: a structural perspective. Curr Opin Struct Biol. 37: 62–70.

Castoldi, M., and A.V. Popov. 2003. Purification of brain tubulin through two cycles of polymerization-depolymerization in a high-molarity buffer. Protein expression and purification. 32: 83–88.

Chowdhury, S., S.A. Ketcham, T.A. Schroer, and G.C. Lander. 2015. Structural organization of the dynein-dynactin complex bound to microtubules. Nat Struct Mol Biol. 22: 345–347.

Cianfrocco, M.A., M.E. DeSantis, A.E. Leschziner, and S.L. Reck-Peterson. 2015. Mechanism and regulation of cytoplasmic dynein. Annu Rev Cell Dev Biol. 31: 83–108.

Derewenda, U., C. Tarricone, W.C. Choi, D.R. Cooper, S. Lukasik, F. Perrina, A. Tripathy, M.H. Kim, D.S. Cafiso, A. Musacchio, and Z.S. Derewenda. 2007. The structure of the coiled-coil domain of Ndel1 and the basis of its interaction with Lis1, the causal protein of Miller-Dieker lissencephaly. Structure. 15: 1467–1481.

DeWitt, M.A., C.A. Cypranowska, F.B. Cleary, V. Belyy, and A. Yildiz. 2015. The AAA3 domain of cytoplasmic dynein acts as a switch to facilitate microtubule release. Nat Struct Mol Biol. 22: 73–80.

Dix, C.I., H.C. Soundararajan, N.S. Dzhindzhev, F. Begum, B. Suter, H. Ohkura, E. Stephens, and S.L. Bullock. 2013. Lissencephaly-1 promotes the recruitment of dynein and dynactin to transported mRNAs. J Cell Biol. 202: 479–494.

Dobyns, W.B., O. Reiner, R. Carrozzo, and D.H. Ledbetter. 1993. Lissencephaly. A brain malformation associated with deletion of the LIS1 gene located at chromosome 17p13. JAMA. 270: 2838–2842.

Duellberg, C., M. Trokter, R. Jha, I. Sen, M.O. Steinmetz, and T. Surrey. 2014. Reconstitution of a hierarchical +TIP interaction network controlling microtubule end tracking of dynein. Nat Cell Biol. 16: 804–811.

Edelstein, A., N. Amodaj, K. Hoover, R. Vale, and N. Stuurman. 2010. Computer control of microscopes using microManager. Curr Protoc Mol Biol. Chapter 14:Unit14 20.

Egan, M.J., K. Tan, and S.L. Reck-Peterson. 2012. Lis1 is an initiation factor for dynein-driven organelle transport. The Journal of cell biology. 197: 971–982.

Feng, Y., E.C. Olson, P.T. Stukenberg, L.A. Flanagan, M.W. Kirschner, and C.A. Walsh. 2000. LIS1 regulates CNS lamination by interacting with mNudE, a central component of the centrosome. Neuron. 28: 665–679.

Gill, S.R.T., T.A. Schroer, I. Szilak, E.R. Steuer, M.P. Sheetz, and D.W. Cleveland. 1991. Dynactin, a conserved, ubiquitously expressed component of an activator of vesicle motility mediated by cytoplasmic dynein. J. Cell Biol. 115: 1639–1650.

Goshima, G., F. Nedelec, and R.D. Vale. 2005. Mechanisms for focusing mitotic spindle poles by minus end-directed motor proteins. J Cell Biol. 171: 229–240.

Heald, R., R. Tournebize, T. Blank, R. Sandaltzopoulos, P. Becker, A. Hyman, and E. Karsenti. 1996. Self-organization of microtubules into bipolar spindles around artificial bipolar spindles around artifical chromosomes in Xenopus egg extracts. Nature. 382: 420–425.

Hoang, H.T., M.A. Schlager, A.P. Carter, and S.L. Bullock. 2017. DYNC1H1 mutations associated with neurological diseases compromise processivity of dynein-dynactin-cargo adaptor complexes. Proc Natl Acad Sci U S A. 114:E1597–E1606.

Huang, J., A.J. Roberts, A.E. Leschziner, and S.L. Reck-Peterson. 2012. Lis1 acts as a “clutch” between the ATPase and microtubule-binding domains of the dynein motor. Cell. 150: 975–986.

Imai, H., T. Shima, K. Sutoh, M.L. Walker, P.J. Knight, T. Kon, and S.A. Burgess. 2015. Direct observation shows superposition and large scale flexibility within cytoplasmic dynein motors moving along microtubules. Nature communications. 6: 8179.

Kardon, J.R., S.L. Reck-Peterson, and R.D. Vale. 2009. Regulation of the processivity and intracellular localization of Saccharomyces cerevisiae dynein by dynactin. Proc Natl Acad Sci U S A. 106: 5669–5674.

Kardon, J.R., and R.D. Vale. 2009. Regulators of the cytoplasmic dynein motor. Nat Rev Mol Cell Biol. 10: 854–865.

King, S.J., and T.A. Schroer. 2000. Dynactin increases the processivity of the cytoplasmic dynein motor. Nat Cell Biol. 2: 20–24.

Kon, T., T. Oyama, R. Shimo-Kon, K. Imamula, T. Shima, K. Sutoh, and G. Kurisu. 2012. The 2.8 A crystal structure of the dynein motor domain. Nature. 484: 345–350.

Levy, J.R., C.J. Sumner, J.P. Caviston, M.K. Tokito, S. Ranganathan, L.A. Ligon, K.E. Wallace, B.H. LaMonte, G.G. Harmison, I. Puls, K.H. Fischbeck, and E.L. Holzbaur. 2006. A motor neuron disease-associated mutation in p150Glued perturbs dynactin function and induces protein aggregation. J Cell Biol. 172: 733745.

Lipka, J., M. Kuijpers, J. Jaworski, and C.C. Hoogenraad. 2013. Mutations in cytoplasmic dynein and its regulators cause malformations of cortical development and neurodegenerative diseases. Biochem Soc Trans. 41: 1605–1612.

Lloyd, T.E., J. Machamer, K. O’Hara, J.H. Kim, S.E. Collins, M.Y. Wong, B. Sahin, W. Imlach, Y. Yang, E.S. Levitan, B.D. McCabe, and A.L. Kolodkin. 2012. The p150(Glued) CAP-Gly domain regulates initiation of retrograde transport at synaptic termini. Neuron. 74: 344–360.

Markus, S.M., and W.L. Lee. 2011. Regulated offloading of cytoplasmic dynein from microtubule plus ends to the cortex. Dev Cell. 20: 639–651.

McKenney, R.J., W. Huynh, M.E. Tanenbaum, G. Bhabha, and R.D. Vale. 2014. Activation of cytoplasmic dynein motility by dynactin-cargo adapter complexes. Science. 345: 337–341.

McKenney, R.J., W. Huynh, R.D. Vale, and M. Sirajuddin. 2016. Tyrosination of alpha-tubulin controls the initiation of processive dynein-dynactin motility. EMBO J.

McKenney, R.J., M. Vershinin, A. Kunwar, R.B. Vallee, and S.P. Gross. 2010. LIS1 and NudE Induce a Persistent Dynein Force-Producing State. Cell. 141: 304–316.

McKenney, R.J., S.J. Weil, J. Scherer, and R.B. Vallee. 2011. Mutually exclusive cytoplasmic dynein regulation by NudE-Lis1 and dynactin. The Journal of biological chemistry. 286: 39615–39622.

Merdes, A., K. Ramyar, J.D. Vechio, and D.W. Cleveland. 1996. A complex of NuMA and cytoplasmic dynein is essential for mitotic spindle assembly. Cell. 87: 447–458.

Moughamian, A.J., G.E. Osborn, J.E. Lazarus, S. Maday, and E.L. Holzbaur. 2013. Ordered recruitment of dynactin to the microtubule plus-end is required for efficient initiation of retrograde axonal transport. J Neurosci. 33: 13190–13203.

Nicholas, M.P., F. Berger, L. Rao, S. Brenner, C. Cho, and A. Gennerich. 2015. Cytoplasmic dynein regulates its attachment to microtubules via nucleotide state-switched mechanosensing at multiple AAA domains. Proc Natl Acad Sci U S A. 112: 6371–6376.

Niethammer, M., D.S. Smith, R. Ayala, J. Peng, J. Ko, M.S. Lee, M. Morabito, and L.H. Tsai. 2000. NUDEL is a novel Cdk5 substrate that associates with LIS1 and cytoplasmic dynein. Neuron. 28: 697–711.

Nirschl, J.J., M.M. Magiera, J.E. Lazarus, C. Janke, and E.L. Holzbaur. 2016. alpha-Tubulin Tyrosination and CLIP-170 Phosphorylation Regulate the Initiation of Dynein-Driven Transport in Neurons. Cell reports. 14: 2637–2652.

Nyarko, A., Y. Song, and E. Barbar. 2012. Intrinsic disorder in dynein intermediate chain modulates its interactions with NudE and dynactin. J Biol Chem. 287: 2488424893.

Olenick, M.A., M. Tokito, M. Boczkowska, R. Dominguez, and E.L. Holzbaur. 2016. Hook Adaptors Induce Unidirectional Processive Motility by Enhancing the Dynein-Dynactin Interaction. J Biol Chem. 291: 18239–18251.

Pandey, J.P., and D.S. Smith. 2011. A Cdk5-dependent switch regulates Lis1/Ndel1/dynein-driven organelle transport in adult axons. J Neurosci. 31: 17207–17219.

Perlson, E., S. Maday, M.M. Fu, A.J. Moughamian, and E.L. Holzbaur. 2010. Retrograde axonal transport: pathways to cell death? Trends in neurosciences. 33: 335–344.

Reddy, B.J., M. Mattson, C.L. Wynne, O. Vadpey, A. Durra, D. Chapman, R.B. Vallee, and S.P. Gross. 2016. Load-induced enhancement of Dynein force production by LIS1-NudE in vivo and in vitro. Nature communications. 7: 12259.

Sasaki, S., A. Shionoya, M. Ishida, M.J. Gambello, J. Yingling, A. Wynshaw-Boris, and S. Hirotsune. 2000. A LIS1/NUDEL/cytoplasmic dynein heavy chain complex in the developing and adult nervous system. Neuron. 28: 681–696.

Schlager, M.A., H.T. Hoang, L. Urnavicius, S.L. Bullock, and A.P. Carter. 2014. In vitro reconstitution of a highly processive recombinant human dynein complex. EMBO J. 33: 1855–1868.

Schmidt, H., R. Zalyte, L. Urnavicius, and A.P. Carter. 2015. Structure of human cytoplasmic dynein-2 primed for its power stroke. Nature. 518: 435–438.

Schroer, T.A., and M.P. Sheetz. 1991. Two activators of microtubule-based vesicle transport. J. Cell Biol. 115: 1309–1318.

Tai, C.Y., D.L. Dujardin, N.E. Faulkner, and R.B. Vallee. 2002. Role of dynein, dynactin, and CLIP-170 interactions in LIS1 kinetochore function. J Cell Biol. 156: 959–968.

Tarricone, C., F. Perrina, S. Monzani, L. Massimiliano, M.H. Kim, Z.S. Derewenda, S. Knapp, L.H. Tsai, and A. Musacchio. 2004. Coupling PAF signaling to dynein regulation: structure of LIS1 in complex with PAF-acetylhydrolase. Neuron. 44: 809–821.

Torisawa, T., A. Nakayama, K. Furuta, M. Yamada, S. Hirotsune, and Y.Y. Toyoshima. 2011. Functional dissection of LIS1 and NDEL1 towards understanding the molecular mechanisms of cytoplasmic dynein regulation. The Journal of biological chemistry. 286: 1959–1965.

Toropova, K., S. Zou, A.J. Roberts, W.B. Redwine, B.S. Goodman, S.L. Reck-Peterson, and A.E. Leschziner. 2014. Lis1 regulates dynein by sterically blocking its mechanochemical cycle. Elife. 3.

Urnavicius, L., K. Zhang, A.G. Diamant, C. Motz, M.A. Schlager, M. Yu, N.A. Patel, C.V. Robinson, and A.P. Carter. 2015. The structure of the dynactin complex and its interaction with dynein. Science. 347: 1441–1446.

Vallee, R.B., R.J. McKenney, and K.M. Ori-McKenney. 2012. Multiple modes of cytoplasmic dynein regulation. Nature Cell Biology. 14: 224–230.

Vallee, R.B., G.E. Seale, and J.W. Tsai. 2009. Emerging roles for myosin II and cytoplasmic dynein in migrating neurons and growth cones. Trends Cell Biol. 19: 347–355.

Vallee, R.B., and J.W. Tsai. 2006. The cellular roles of the lissencephaly gene LIS1, and what they tell us about brain development. Genes Dev. 20: 1384–1393.

Yi, J.Y., K.M. Ori-McKenney, R.J. McKenney, M. Vershinin, S.P. Gross, and R.B. Vallee. 2011. High-resolution imaging reveals indirect coordination of opposite motors and a role for LIS1 in high-load axonal transport. The Journal of cell biology. 195: 193–201.

Zylkiewicz, E., M. Kijanska, W.C. Choi, U. Derewenda, Z.S. Derewenda, and P.T. Stukenberg. 2011. The N-terminal coiled-coil of Ndel1 is a regulated scaffold that recruits LIS1 to dynein. J Cell Biol. 192:433–445.

